# Light Impact on Thalamo-Cortical Connectivity During a Cognitive Task Depends on Time of Day and is Different in Teenagers

**DOI:** 10.1101/2025.04.03.647005

**Authors:** Roya Sharifpour, Islay Campbell, Ilenia Paparella, Fermin Balda Aizpurua, Elise Beckers, Nasrin Mortazavi, Ekaterina Koshmanova, John Read, Fabienne Collette, Christophe Phillips, Puneet Talwar, Laurent Lamalle, Mikhail Zubkov, Gilles Vandewalle

**Author notes:** Corresponding author: Gilles Vandewalle, GIGA-Cyclotron Research Centre-Human Imaging, Bâtiment B30, 8 Allée du Six Août, University of Liège-Sart Tilman, 4000 Liège, Belgium. Department of Translational Neuroimaging, Hôpital Erasme, Hôpital Universitaire de Bruxelles, Université libre de Bruxelles, Brussels, Belgium (Roya Sharifpour). Sir Jules Thorn Sleep and Circadian Neuroscience Institute, Nuffield Department of Clinical Neurosciences, University of Oxford, Oxford, UK.

## Abstract

**Background:** Light affects not only vision but also attention, alertness, and cognition, primarily through intrinsically photosensitive retinal ganglion cells sensitive to blue-wavelength light. While previous research has shown that light influences brain regional activity, its impact on brain connectivity remains unclear.

**Methods:** Using 7-Tesla fMRI, this study examined how light illuminance modulates canonical left-lateralized verbal thalamo-prefrontal-parietal network connectivity during an auditory executive task. Fifty-five participants, including young adults (19–30 y, 23 Females: scanned in the morning or evening) and adolescents (15–18 y, 6 Females: scanned only in the evening), were studied.

**Results:** We first found that across all the groups, moderate blue-enriched light strengthened parietal-to-frontal connectivity, while low-illuminance orange light enhanced thalamus-to-parietal connectivity. When focusing on time-of-day differences in young adults, we found that higher-illuminance blue-enriched light strengthened thalamus-to-prefrontal connectivity in the morning compared to the evening. When focusing on developmental stages differences, during the evening session only, we observe that moderate blue-enriched light enhanced thalamus-to-parietal connectivity in adolescents compared to young adults.

**Conclusion:** These findings reinforce the view that the thalamus plays a key role in mediating the impact of light on cognition and the impact of light depends on the context of light administration, i.e. with timing and age.

## Introduction

Light influences not only the visual system but also non-visual functions, such as physiological, hormonal, and neurobehavioral responses[1–3]. These non-image forming (NIF) effects are primarily mediated by intrinsically photosensitive retinal ganglion cells (ipRGCs), a third type of photoreceptor in the retina, on top of rods and cones, that are most sensitive to short-wavelength (blue) light at approximately 480 nm[4]. IpRGCs directly project to a wide range of brain structures, including the hypothalamus, thalamus, and amygdala (see, e.g., ref[1] for a review of these projections). Through these widespread projections, light exposure has direct and indirect effects on the regulation of circadian rhythms, sleep‒wake cycles and mood, as well as on emotions, attention, alertness and cognitive performance[5–7]. The acute NIF effects of light on brain function are modulated by several factors, including time of day, with evidence from human brain studies suggesting stronger effects in the morning than in the evening[8]. These effects may also vary across lifespan[9,10]. In particular, research indicates that adolescents may respond more strongly to light than adults do[9].

Several functional magnetic resonance imaging (fMRI) studies showed that light modulate blood oxygen level-dependent (BOLD) signals of the brain including in key subcortical regions for sleep and wakefulness regulation such as the hypothalamus and thalamus. These studies further demonstrated that light interacted with ongoing cognition such that several cortical regions involved in the ongoing non-visual cognitive process saw their activity increased during or following exposure to light, thereby influencing cognitive performance[7,11–13] (for a comprehensive review, see ref[1]). However, the neural mechanisms and modulating factors by which light affects cognition are not yet fully understood. We previously hypothesized that the influence of light on cognition might be mediated by changes in brain connectivity, particularly from subcortical structures to the cortical regions engaged in the ongoing cognitive task. The thalamus, a crucial subcortical relay center that notably interfaces between arousal and cognition[14], is proposed to play an essential role in how light influences information processing in the brain during non-visual cognitive tasks, as evidenced by fMRI studies showing thalamic activation in response to light during cognitive performance[12,13,15]. In support of our hypotheses, we recently showed that during an attentional task, light illuminance influences the connectivity from the thalamus (pulvinar) to the intraparietal sulcus (IPS), key regions involved in attentional regulation[16]. Whether and how these observations extend to other cognitive domains is not known.

Here, we used 7T fMRI to examine how light influences the effective connectivity within a canonical left-lateralized three-region verbal network that plays crucial roles in executive functions, comprising the thalamus, over the mediodorsal nucleus (MDN), the parietal cortex, over the supramarginal gyrus (SMG) in the immediate vicinity of the IPS, and the prefrontal cortex, over the inferior frontal junction (IFJ), next to the middle frontal gyrus (MFG)[17–19]. We also examined how different times of the day and distinct age groups, particularly adolescents and young adults, affect the connectivity among these three regions. This was done first by comparing connectivity in the morning versus in the evening sessions in young adults only, and second by assessing the connectivity in young adults and teenagers in the evening only, as it may constate a critical time for the impact of light in teenagers[20]. We found that blue-enriched light commonly influenced the network at both times of day and in both age groups. Light impact on the connectivity of the network further depended on time of day, with larger impact on the thalamus-to-prefrontal connectivity in the morning (in young adult), and on development stage with a larger impact on thalamus-to-parietal connectivity in teenagers (in the evening). Our findings support the hypothesis that blue-wavelength light impacts non-visual cognitive functions by modulating task-dependent information flow, particularly from subcortical to cortical regions and unravel part of the factors modulating light interaction with cognition.

## Materials and Methods

This study is part of a larger study that has resulted in several publications using different participant subsets[16,21–23].

### Participants

Between February 2021 and September 2023, 55 healthy volunteers aged 15--30 years (22.0±4.6 y, 29 females), including 18 adolescents (16.7±1.1 y, 6 females) and 37 young adults (24.6±3.3 y, 23 females), participated in the study. The exclusion criteria included a body mass index (BMI) >28; recent psychiatric history; severe trauma; sleep disorders; addiction; chronic medication; smoking; excessive alcohol consumption (>14 units per week) or caffeinated drinks (>4 cups per day); night shift work within the past year; transmeridian travel in the past 2 months; and a history of ophthalmic disorders. The participants also completed questionnaires assessing anxiety (21-item Beck Anxiety Inventory)[24], mood (21-item Beck Depression Inventory-II)[25], sleep quality (Pittsburgh Sleep Quality Index)[26], daytime sleepiness (Epworth Sleepiness Scale)[27], insomnia (Insomnia Severity Index)[28], chronotype (Horne-Östberg)[29], and seasonal changes in mood and behavior (Seasonal Pattern Assessment Questionnaire)[30]. **Table 1** summarizes the demographic characteristics of the participants. We stress that all three groups are of similar size and therefore relatively well balanced to seek our effect of interest. We further note that women represent between 50 and 65% of each group meaning that sex effects can be adequately controlled for by including a sex covariate in the statistical analyses. The analyses are, however, most likely not powerful enough to detect sex difference in our effects of interest.

**Table 1.** Demographic characteristics of the participants included in the analyses.

|  | Adults_Evening<br>(AE) | Adults_Morning<br>(AM) | Teenagers<br>(T) | Comparison (t-test) |  |
| --- | --- | --- | --- | --- | --- |
|  |  |  |  | AE vs AM | AE vs T |
| Number of Subjects | 17 | 20 | 18 |  |  |
| Age (mean±SD) | 25.2±4.0 | 24.1±2.5 | 16.7±1.1 | P=0.30 | **P<0.0001** |
| Sex: Female (Male) | 10(7) | 13(7) | 6(12) | P=0.71 | P=0.14 |
| Body mass index (kg/m<sup>2</sup>) | 22.1±2.0 | 21.3±2.3 | 21.5±2.5 | P=0.30 | P=0.42 |
| Depression level (BDI-II<sup>a</sup>) | 6.6±3.5 | 6.2±5.5 | 5.6±4.9 | P=0.79 | P=0.56 |
| Anxiety Level (BAI<sup>b</sup>) | 5.1±3.1 | 5.3±6.1 | 6.1±6.8 | P=0.92 | P=0.77 |
| Chronotype (HO<sup>c</sup>) | 43.1±7.3 | 51.4±7.6 | 42.8±7.6 | **P=0.003** | P=0.91 |
| Subjective Sleep Quality<br>(PSQI<sup>d</sup>) | 4.1±1.9 | 3.7±2.7 | 4.3±1.6 | P=0.63 | P=0.77 |
| Habitual daytime Sleepiness<br>(ESS<sup>e</sup>) | 7.0±2.4 | 5.9±3.0 | 6.0±4.2 | P=0.33 | P=0.50 |
| Insomnia symptoms (ISI<sup>f</sup>) | 6.2±4.9 | 4.4±3.4 | 4.2±5.7 | P=0.22 | P=0.79 |
| Season of experiment * | -0.43±0.51 | -0.14±0.67 | -0.36±0.58 | P=0.14 | P=0.72 |
| Seasonality (SPAQ<sup>g</sup>) | 1.3±0.9 | 0.9±0.7 | 0.7±1.0 | P=0.26 | P=0.16 |

<sup>a</sup>. Beck's Depression Inventory II
<sup>b</sup>. Beck's Anxiety Inventory
<sup>c</sup>. Horne and Östberg Questionnaire
<sup>d</sup>. Pittsburg Sleep Quality Index
<sup>e</sup>. EPWORTH Sleepiness Scale
<sup>f</sup>. Insomnia Severity Index
<sup>g</sup>. Seasonal Pattern Assessment Questionnaire
See the text for reference of the questionnaires.
\* Cosine (acquisition day of year\*360/365); 21 December=0°
\*\* All included participants were right-handed.
Bold values indicate statistical significance at the p<0.05 level.

### Protocol and light exposure

The participants followed a loose sleep‒wake schedule for seven days before the in-lab experiment [±1 h; verified with actigraphy] to prevent excessive sleep deprivation while maintaining realistic conditions. Depending on their fMRI schedule (morning or evening), they arrived at the laboratory either 1.5 hours after waking up or 2 hours before bedtime. Among adults, 20 out of 37 participated in the morning fMRI session, whereas the remaining adults completed the evening session. Due to the general concerns regarding evening light consumption during adolescence, all teenagers completed their fMRI sessions in the evening (we further anticipated that scholar constraints would prevent us from completing fMRI sessions in the morning).

To control for previous effects of short-term prior light exposure, upon arrival, participants first underwent 5 minutes of exposure to a relatively bright polychromatic white light (∼1000 lux), followed by 45 minutes of dim light (< 10 lux). During this period, they received instructions for the fMRI study and practiced the executive task (n-back task) on a laptop (**Figure 1-A**). While the bright polychromatic light may have influenced melatonin levels, existing literature suggests that any residual effects would likely have dissipated by the time of measurement[31,32].

**Figure 1:**
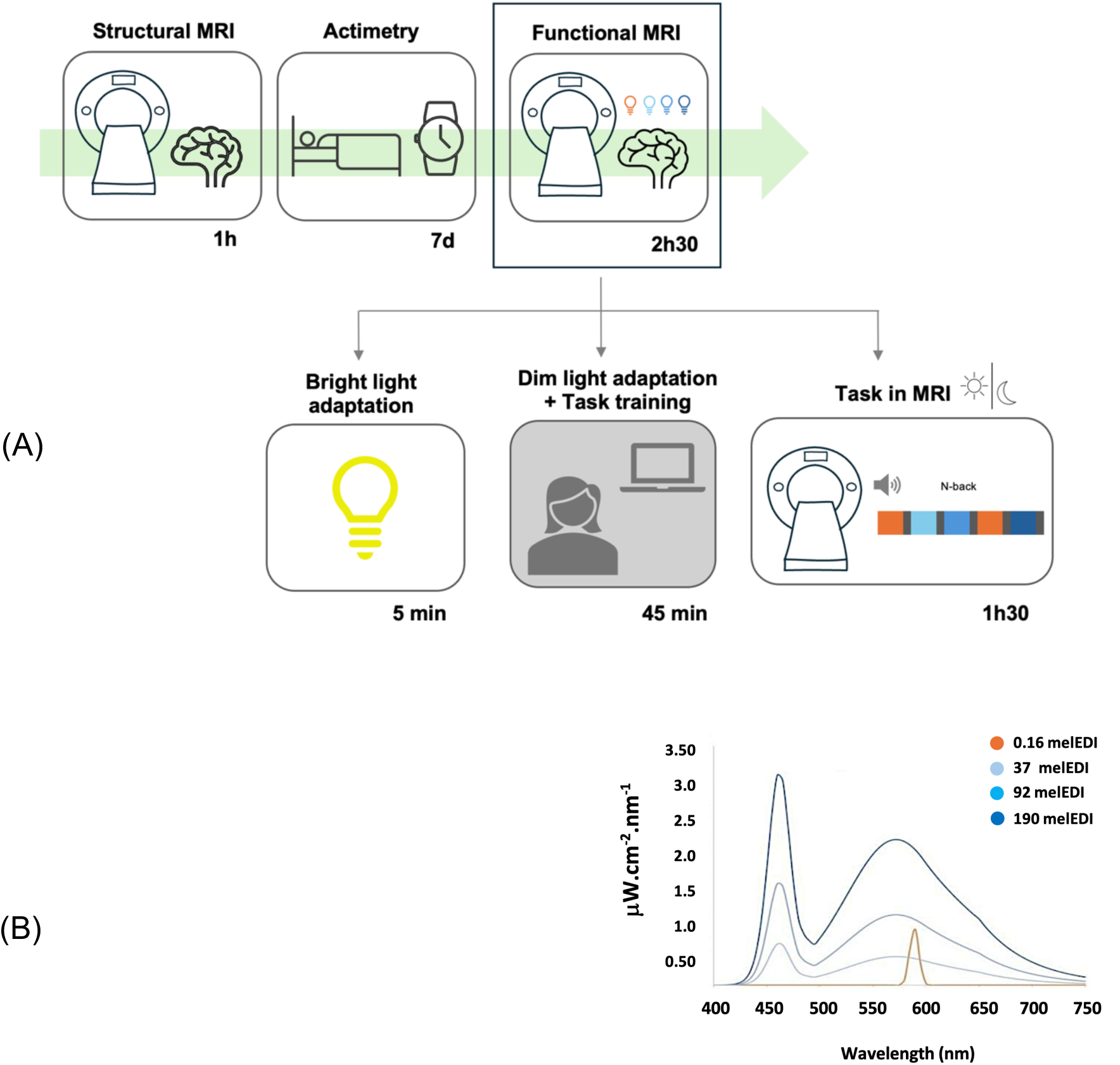
Graphical representation of the experimental protocol. **(A)**: Participants completed a structural MRI session followed by a functional MRI scan 7 days later, with sleep‒wake patterns monitored in between. Before the functional MRI scan, the participants were exposed to bright white light for 5 minutes, followed by a training session for auditory tasks inside the scanner for 45 minutes in darkness (<10 lux). During the fMRI, they performed a working memory task (N-back) while being exposed to alternating blocks of blue-enriched or orange monochromatic light, separated by darkness periods. **(B):** Spectrum of the 4 light conditions to which participants were exposed during the N-back task. The duration, in days, hours or minutes, of each step is provided below each box. The full light characteristics are shown in **Table S1**.

Inside the scanner, participants performed an auditory letter variant of the n-back task at two difficulty levels, 0-back and 2-back, and responded to the task via an MRI-compatible response box. This task involves maintaining and continuously updating relevant information in working memory. In the 0-back task, which is less demanding and serves as a control for baseline brain activity, participants responded whenever the current letter matched a predefined letter. In the more challenging 2-back task, participants determined if the current letter was identical to the one presented two stimuli earlier. We used a block design with stimuli presented in 38 blocks, each lasting approximately 33 seconds, comprising 19 blocks of the 2-back task. The order of the 0-back and 2-back tasks was pseudorandomized over the entire session.

The participants were alternately maintained in darkness (<0.01 lux) or exposed to blue-enriched cool polychromatic light (6500K) at varying illuminance levels (37, 92, 190 melanopic equivalent daylight illuminance (mel-EDI) lux) or a control monochromatic orange light (590 nm; 0.16 mel-EDI lux), to which ipRGCs are nearly insensitive (**Figure 1-B**). The blue-enriched light illuminances were set according to the technical characteristics of the light source and to keep the overall photon flux similar to that in prior 3T fMRI studies by our team using the same task (between ∼10^12^ and 10^14^ ph/cm²/s) [15,33,34]. Orange light was introduced as a control for visual stimulation for potential secondary whole-brain analyses. For the present analyses, we discarded color differences between the light conditions and only considered illuminance as indexed by mel-EDI lux, constituting a limitation of our study. The detailed methodology for the light setup and its delivery into the MRI scanner can be found in our previous publication[16]. There were 8 blocks per light condition and 6 blocks for the darkness condition. There were also darkness periods (∼10 s, <0.01 lux) without tasks between blocks with different light conditions to ensure that the light exposure in one block did not affect brain activity in the subsequent block. During all fMRI sessions, pupil size was recorded simultaneously with an eye-tracking device (EyeLink 1000Plus, SR-Research, Ottawa Canada). Eye tracking confirmed that participants kept their eyes open during the scan. We previously reported that pupil constriction was stronger under higher mel-EDI levels, based on data from a subset of young adults scanned in the morning. This subset partially overlaps with the morning young adult group in the present study[23]. However, due to equipment-related data loss, the amount of usable pupillometry data in the evening young adult group and in teenagers was insufficient to support reliable analysis in this study. The entire experiment was designed via OpenSesame (3.2.8)[35].

### MRI Data Acquisition

Structural and functional MRI data were acquired using a MAGNETOM Terra 7 Tesla MRI system (Siemens Healthineers, Erlangen, Germany) with a one-channel transmit and a 32-channel receive head coil (Nova Medical, MA, USA). To reduce dielectric artifacts, dielectric pads (Multiwave imaging, Marseille, France) were placed between participants’ heads and the receiver coil. Multi-band Gradient-Recalled Echo - Echo-Planar Imaging (GRE-EPI) sequence was used for acquiring multislice T2*-weighted functional images, with axial slice orientation and specific parameters, including TR =2340 ms, TE =24 ms, FA =90°, no interslice gap, field of view (FoV) = 224 mm × 224 mm, matrix size = 160 × 160 × 86, and voxel size of (1.4 × 1.4 × 1.4) mm^3^. The initial three scans were excluded to mitigate saturation effects. Additionally, for anatomical reference, a high-resolution T1-weighted image was acquired via a Magnetization-Prepared with 2 RApid Gradient Echoes (MP2RAGE) sequence, with specific parameters including TR =4300 ms, TE =1.98 ms, FA =5°/6°, TI =940 ms/2830 ms, bandwidth = 240 Hz, matrix size =256×256×224, acceleration factor =3, and voxel size = 0.75×0.75×0.75 mm^3^.

### Preprocessing

MP2RAGE images were processed using a statistical parametric mapping (SPM12) extension, which relies on a regularization factor to limit background noise[36]. These denoised images were subsequently automatically reoriented using SPM and corrected for intensity bias caused by field inhomogeneity using the bias correction method within the SPM’s “unified segmentation” approach[37]. Brain extraction was performed on the denoised, reoriented, and bias-corrected images to avoid potential co-registration issues arising from the use of dielectric pads during the scans. This process was performed using SynthStrip[38].

The fMRI time series underwent estimation of static and dynamic susceptibility-induced variance using voxel-displacement maps computed from phase and magnitude images. “Realign & Unwarp” was then applied to the EPI images to correct for head motion and for static and dynamic susceptibility-induced variance. Realigned and distortion-corrected EPI images were then subjected to brain extraction with SynthStrip, followed by smoothing using a Gaussian kernel with a full width at half maximum (FWHM) of 3 mm. First-level analysis for each subject was conducted in their native space to prevent potential errors introduced by co-registration. Prior to second-level analysis, contrast maps from first-level analyses were transferred to the subject structural space, then to the group template space, and ultimately to the MNI space (1×1×1 mm³), with all performed using Advanced Normalization Tools (ANTs; Penn Image Computing and Science Laboratory, UPenn, USA; https://stnava.github.io/ANTs/).

### Univariate Analysis

For each subject, changes in brain regional BOLD signals were estimated using a general linear model (GLM). Within the GLM design matrix, both levels of the task (0-back and 2-back) were modeled as the main regressors, and light was included as a modulatory regressor, reflecting the varying levels of light illuminance measured in mel-EDI lux. These regressors were then convolved with the canonical hemodynamic response function to generate the predicted BOLD response. Movement parameters, as well as cardiac and respiratory parameters derived using the PhysIO Toolbox (Translational Neuromodeling Unit, ETH Zurich, Switzerland), were included as regressors of no interest in the GLM. To eliminate low-frequency drifts, high-pass filtering was applied with a cutoff frequency of 256 Hz. Our contrast of interest focused on regions showing increased activation in response to the 2-back task compared with the 0-back task, independent of the light condition. These contrast images (in MNI space) were then taken for the second-level analysis, employing a random-effects model to examine group-level effects. To control for multiple comparisons, the results were corrected at the voxel level using a family-wise error (FWE) procedure, with a significance threshold set at 0.05.

Based on the group-level results, group peak coordinates of regions of interest (ROIs) in the left hemisphere were identified: the MDN of the thalamus, the SMG, next to the IPS, and the IFJ, next to the MFG (**Figure 3-A**). The left hemisphere was selected because research has demonstrated hemispheric asymmetry in working memory, with the left hemisphere showing greater activation in verbal working memory and the right hemisphere in spatial working memory[39].

These ROIs were selected because the prefrontal and parietal cortices are integral to the phonological loops involved in verbal tasks and are typically engaged during the n-back task[40–42]. Furthermore, thalamus involvement is commonly reported in the context of working memory[43,44]. Additionally, light interacts with the ongoing task by modulating cortical activity, likely via subcortical structures, with the thalamus potentially serving as a central hub. Therefore, we specifically targeted a thalamo–frontal–parietal network. Individual ROIs were then defined by selecting the first activated cluster within a sphere of a specific radius, which was determined based on the size of each nucleus and centered on the group peak coordinates of the SMG (8 mm), IFJ (8 mm), and MDN (5 mm). For the effective connectivity analysis, BOLD time series were used to infer the underlying neural activity. The first principal components (eigenvariates) of the BOLD signal time series within those ROIs were extracted from the individual statistical map, which was thresholded at p = 0.05 uncorrected. For the eigenvariate extraction, the “adjusted” time series was used, which represents the time series after regressing out effects of no interest, via the approach outlined by Zeidman *et al*.[45].

Optimal sensitivity and power analyses in MRI remain under investigation (e.g.[46]) particularly when including an effective connectivity approach. We nevertheless performed a prior sensitivity analysis to get an indication of the minimum detectable effect size in our main analyses, given our sample size. According to G*Power 3 (version 3.1.9.4; [47]), taking into account a power of 0.8, an error rate α of 0.05, a sample of 55 allowed us to detect large effect sizes *r*>0.35 (two-sided; absolute values; CI: .09–.56; R²>.12, R² CI: .008–.31) within a multiple linear regression framework including one tested predictor (illuminance effect) and three covariates (age group/time of day, sex, and BMI).

### Effective Connectivity Analysis and Statistics

We employed dynamic causal modeling (DCM) framework[45], implemented in SPM12, which is a method designed to examine network connectivity and how it is modulated. In our study, DCM was used to investigate whether light modulated task-related neural activity by altering the connectivity between three predefined network ROIs involved in the ongoing task. In the DCM analysis, six inputs were defined within a design matrix and then imported into the DCM framework. These inputs comprised all 0-back and 2-back trials as two separate driving inputs, as well as blocks representing the four light conditions (including the control orange light and blue-enriched cool light at three different mel-EDI levels), each serving as separate modulatory inputs. The DCM model included all intrinsic connectivity among the three regions, along with self-feedback gain control connectivity. Additionally, the model considered the influence of the task on all regions and allowed for the potential modulation of connectivity between regions by all light conditions (**Figure 4-A**).

Time series extracted from individual ROIs were subjected to first-level DCM analysis, where the model was estimated for each subject. We subsequently performed a parametric empirical Bayes (PEB) analysis[48,49] over the first-level DCM parameter estimates. PEB is a hierarchical Bayesian model that evaluates commonalities and differences among subjects in the effective connectivity domain at the group level. This method considers variability in individual connectivity strengths and reduces the influence of subjects with noisy data. Separate PEB analyses were conducted for each matrix (A: intrinsic connectivity, B: modulatory effects and C: effects of driving inputs) to prevent the dilution of evidence by reducing the search space. After the full model (with all connectivity of interest) for each subject was estimated, the PEB approach included Bayesian model reduction (BMR) and averaging (BMA) of the parameters across models weighted by the evidence of each model. Since there is no concept of significance in Bayesian analysis, we have reported only parameters contributing to the model with at least positive evidence, i.e., posterior probability (Pp) exceeding 0.73 (according to ref[50], positive Pp: 0.73–0.95, strong Pp: 0.95–0.99, and very strong Pp: > 0.99).

A generalized linear mixed model (GLMM) was first applied using SAS 9.4 (SAS Institute, NC, USA) to assess the impact of light conditions on task performance, irrespective of connectivity. In this model, task accuracy was the dependent variable, with subject (intercept and slope) effects included as a random factor. Light conditions (five illuminance levels) were treated as repeated measures with an autoregressive correlation structure of type 1 (AR(1)). The fixed effects included the following predictors: light illuminance, age group, time of day, and covariates, including sex and BMI.

A second set of GLMMs was used to determine whether light-modulated connectivity influenced task performance. In these models, task accuracy under the specific lighting condition that modulated connectivity was the dependent variable, with subject (intercept and slope) effects as a random factor. The independent variables included modulated connectivity, age group, and/or time of day, depending on the specific connectivity being analyzed. Sex and BMI were included as covariates of no interest.

All the models were adjusted to account for the distribution of the dependent variables. Tukey adjustment was applied to direct post hoc tests of the primary analysis to correct for multiple comparisons. Participants whose task accuracy was less than 65% were excluded from the behavioral analyses (2 individuals).

## Results

Fifty-five healthy participants underwent fMRI scans either in the morning or evening. These were distributed as follows: 20 young adults scanned in the morning, 17 young adults scanned in the evening, and 18 adolescents scanned in the evening. During the scan, the participants performed an auditory working memory task (n-back) under 4 different light conditions, including low illuminance orange light (0.16 mel-EDI) and three intensities of blue-enriched cool light (37, 92 and 190 mel-EDI) (**Figure 1**).

Performance to the 2-back task was good for all participants (mean±SD of accuracy on the 2-back task: 88.2%±9.0% across all subjects; 88.0%±11.0% across adults with morning fMRI; 90.5%±7.5% across adults with evening fMRI; and 87.1%±8.2% across adolescents). Importantly, light affected performance in the 2-back task across all groups (F(4,208) = 5.55; **P= 0.0003**), with improved accuracy under high illuminance (0.16 & 37 mel-EDI lux < 92 & 190 mel-EDI lux; t ≤ −2.8; P < 0.04) (**Figure 2**), whereas there were no differences between age groups and times of day in the impact of light (F(1,48_time-of-day_/48 _age group_) < 0.05; P > 0.9).

**Figure 2:**
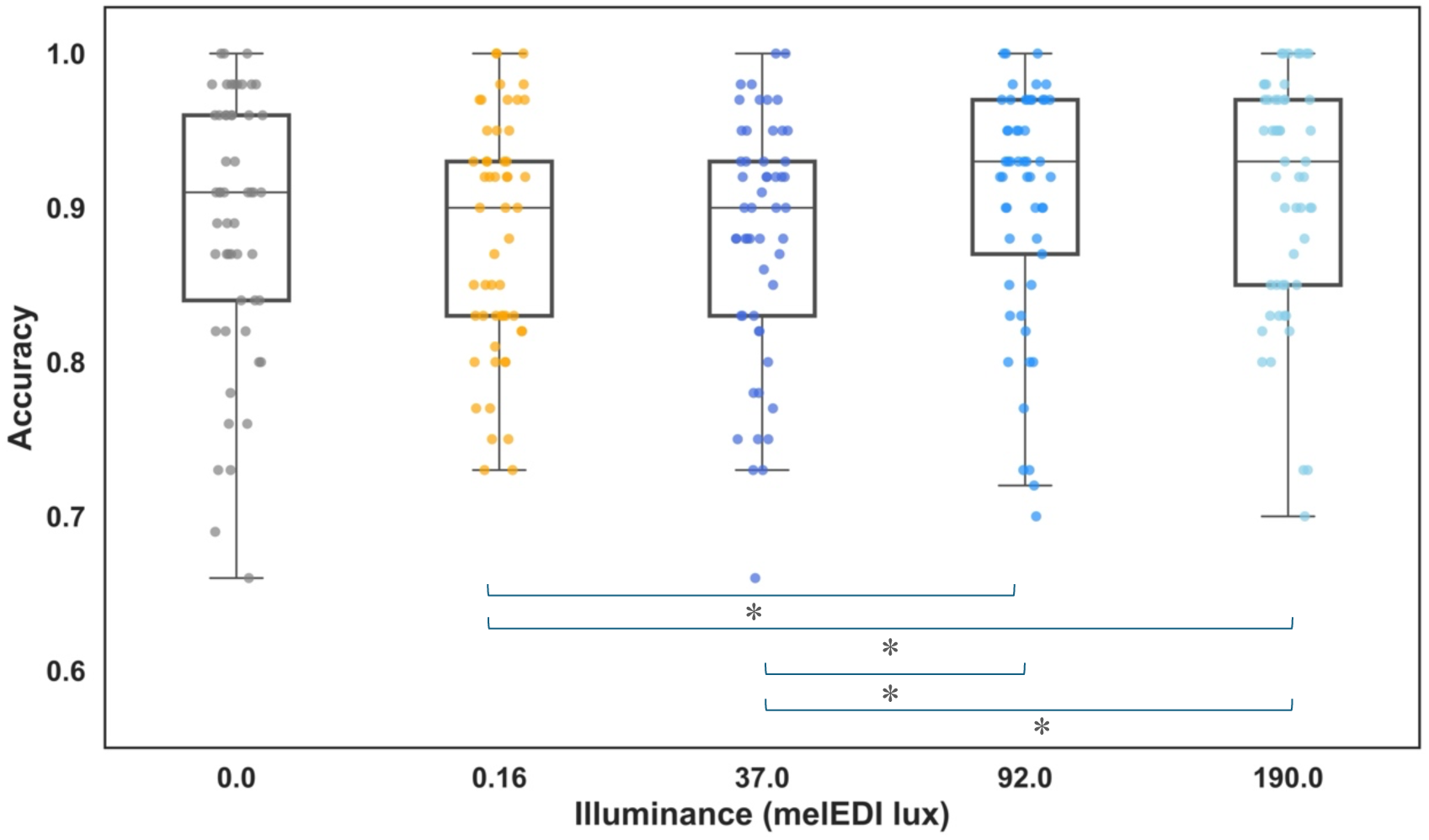
Performance in the 2-back task under each light condition across all groups. (main effect of illuminance: p= 0.0003). Compared with the control orange light (0.16 mel-EDI) and lowest intensity (37 mel-EDI) blue-enriched light, the participants were more accurate in the 2-back task under moderate (92 mel-EDI) and high (190 mel-EDI) blue-enriched lights. * p < 0.05 (post hoc contrasts)

At the brain level, following a standard univariate analysis, we examined how varying illuminance influenced the connectivity between three primary brain regions of interest involved in the task and assessed separately whether the time of day and age affected the impact of light on connectivity.

### Univariate Analysis of the Response to the 2-back vs. 0-back Tasks

After controlling for baseline brain activity via the 0-back task, we conducted a standard univariate analysis to isolate regions of interest that were responsive to the task, irrespective of the light condition (and the age group and time of day). A widespread set of regions showed activation in response to the task, which was consistent with the findings of previous studies (**Table S2**). This analysis confirmed that the task was successful in triggering activation within the thalamus, which is commonly reported in the context of working memory[43,44], and in the prefrontal and parietal cortices, which are typically involved in the verbal n-back task[40–42]. We selected our ROIs as the maximal activation within these areas over the left hemisphere given the verbal nature of the task. Specifically, we observed bilateral activation in the thalamus, the SMG and the MFG, with spatially broader activation in the left hemisphere (which is again in line with the verbal nature of the task; **Figure 3-A**), as well as in the anterior insula and cerebellum (**Figure S1**). Thus, the first principal component of the BOLD time series was extracted from the left MDN, left SMG, and left IFJ, all of which are engaged in ongoing executive processes, to infer their respective neuronal activities. We then used the DCM to examine the effective connectivity among these three regions and determine how light interacted (modulated) the network in each group of subjects.

**Figure 3.**
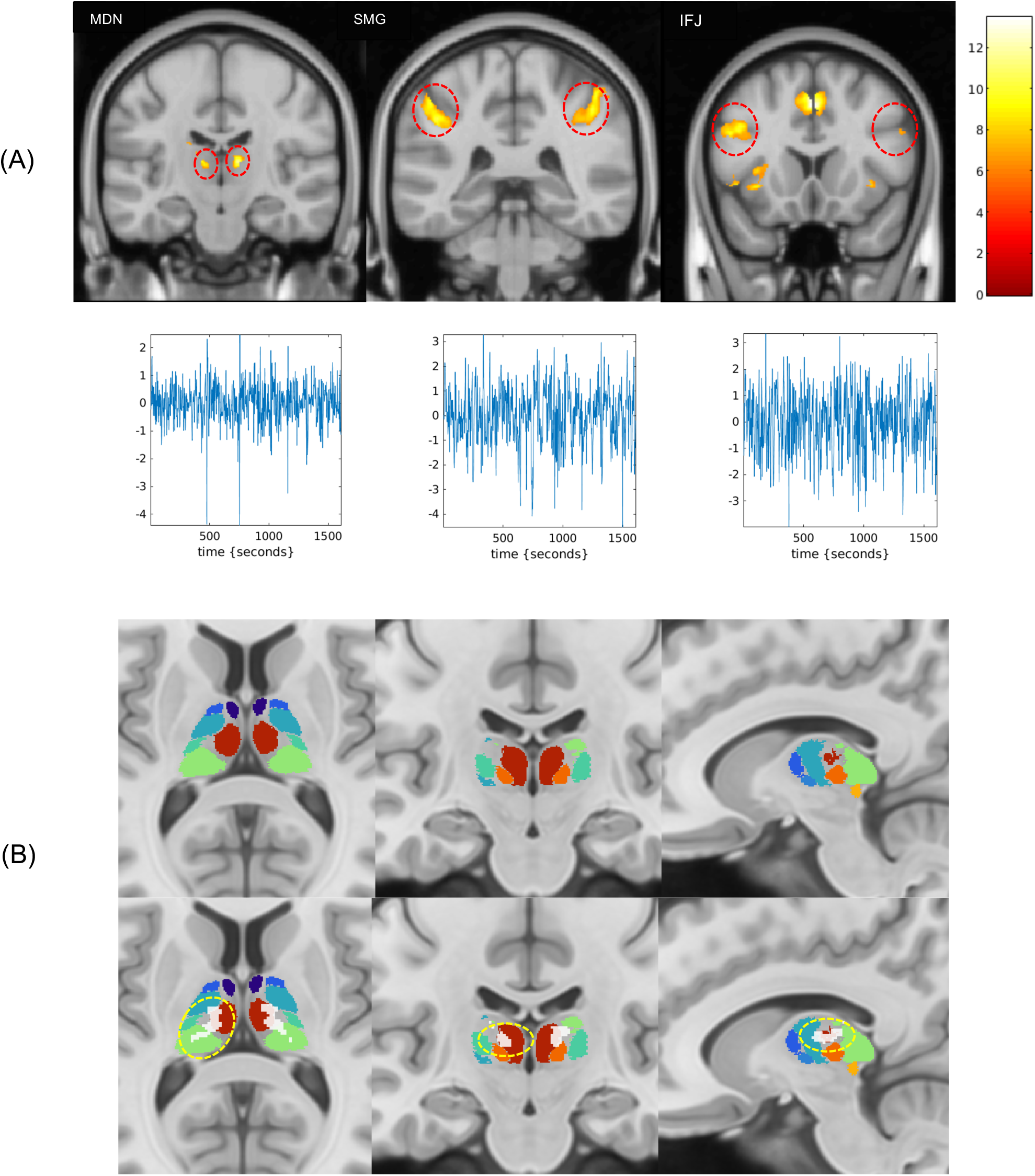
Activation and eigenvariate extracted from the 3 ROIs included in the DCM analysis. **(A)** Left: Bilateral thalamic activation in the mediodorsal nucleus (MDN), Middle: Bilateral parietal activation in the supramarginal gyrus (SMG), Right: Bilateral activation in the inferior frontal junction (IFJ). **(B)** Top row: Thalamus parcellation map[51]: Maroon: MDN, Orange: Centromedian nucleus (CMN), Light Green: Pulvinar, Yellow: Medial geniculate nucleus (MGN), Turquoise: ventral posterolateral (VPL), Dark Turquoise: ventral lateral posterior (VLp), Royal Blue: Ventral anterior nucleus (VAN), Navy: Anteroventral nucleus (AVN). Habenula, lateral geniculate and ventral lateral anterior nuclei are not visible in the sections shown here. Bottom row: Thalamic activity overlaid (shown in white) on the parcellation.

### Connectivity Common to the Three Groups

To compare the modulatory impact of light on connectivity between the morning and evening, as well as between adults and adolescents, we performed a two separate group comparison as part of the same DCM analysis. In that construct, the adult group that underwent fMRI scans in the evening was considered the reference for analyses, i.e., the group common to all comparisons that would provide the light-induced modulation common to all 3 groups. The other two groups, which consisted of young adults scanned in the morning and of adolescents scanned in the evening, were compared against this reference group to identify additive and separate effects related to time of day (for adults) and age differences (in the evening), respectively.

DCM analysis in the reference group (**Figure 4-B**) revealed that, compared with the 0-back task, the 2-back task, a working memory challenge, predominantly activated the IFJ and SMG (Pp=1.0), indicating that these regions are primarily responsible for handling the cognitive demands of the task within our 3-region network. Interestingly, this means that, according to our analyses, the MDN did not receive direct task input within our network, suggesting that its role may be more about facilitating interregional communication rather than direct task-related processing. DCM also yielded very strong evidence (Pp=1.0) for self-inhibition in both the SMG and the MDN, suggesting intrinsic regulatory mechanisms that maintain stable activity levels. DCM further yielded strong evidence for excitatory connectivity from the MDN to both the SMG (Pp=1.0) and the IFJ (Pp=1.0), highlighting that the MDN significantly influences SMG and IFJ activity. In addition, DCM provided strong evidence for bilateral connectivity between the MDN and IFJ (Pp=1.0), indicating a reciprocal relationship, where both regions influence each other with the MDN exerting excitatory effects, whereas the IFJ provides inhibitory feedback. The last intrinsic connectivity with strong evidence was directed inhibitory connectivity from the IFJ to the SMG (Pp=1.0), which highlights the role of the IFJ in driving SMG activity. The overall pattern of excitatory thalamic inputs and inhibitory IFJ inputs supports complex task performance by balancing activation and regulation across these regions.

**Figure 4.**
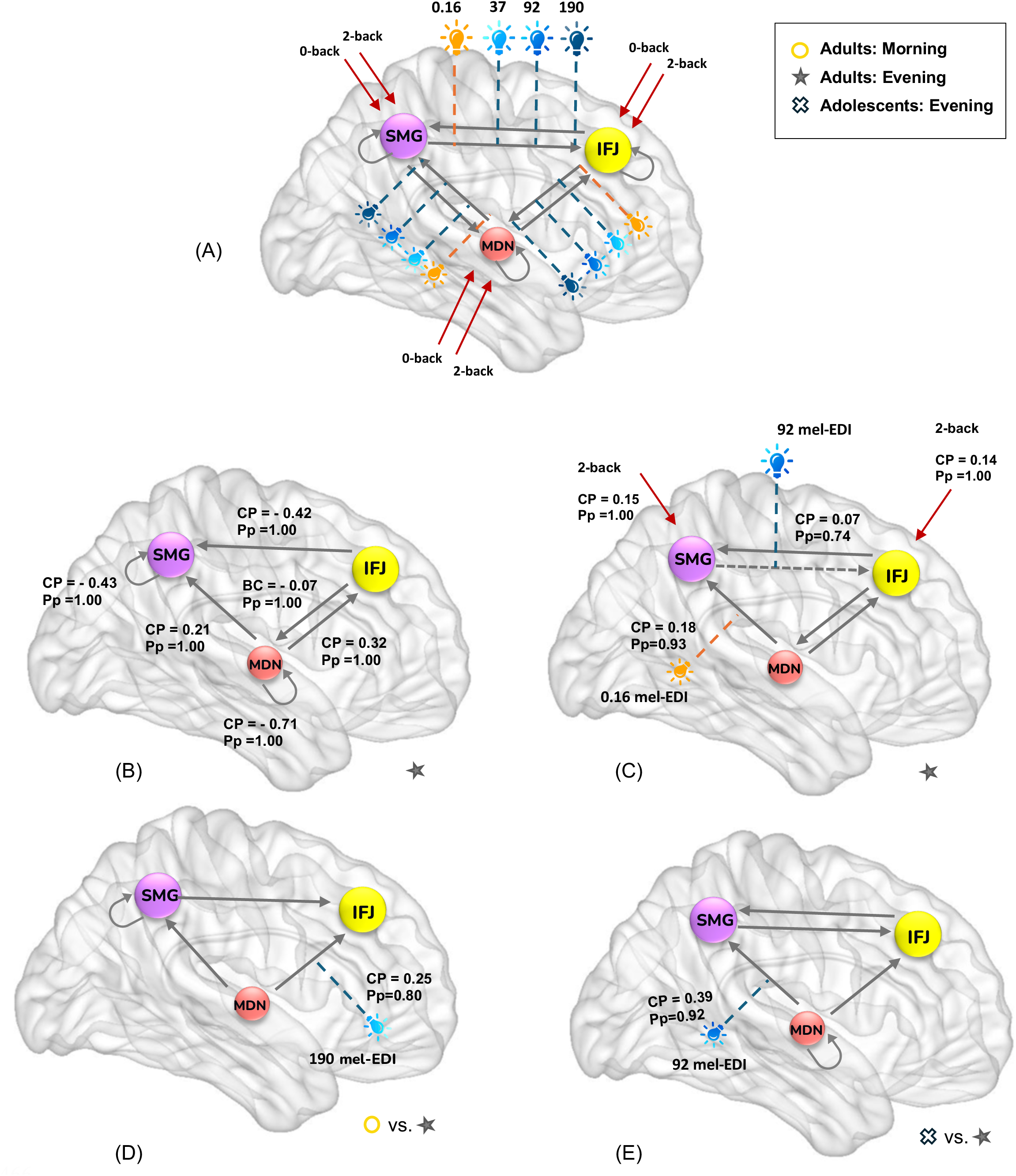
Effective connectivity results: **(A)** initial model tested using the DCM framework. **(B)** Baseline (intrinsic) connectivity parameter (CP) common to all groups, as isolated in the reference group (i.e., Adults with fMRI in the evening). **(C)** Modulatory impact of light conditions on connectivity along with task input in the reference group (common to all groups). **(D)** <u>Differences</u> between the connectivity parameters in the morning compared with those in the evening (in adults). **(E)** <u>Differences</u> in the connectivity parameters between adolescents and adults (both in the evening). IFJ: inferior frontal junction; MDN: mediodorsal nucleus of the thalamus; mel-EDI: melanopic equivalent daytime illuminance. Pp: posterior probability; SMG: supramarginal gyrus; gray arrows: between regions and self-inhibitory connectivity; red arrows: task inputs; colored light bulb: light conditions as connectivity modulators (orange: 0.16 mel-EDI; dark blue: 37 mel-EDI; medium blue: 92 mel-EDI; light blue: 190 mel-EDI).

The primary objective of this study was to evaluate how light modulates connectivity. DCM analysis in the reference group (adults, evening) revealed positive evidence (P=0.74) for a strengthening impact of moderate illuminance (92 mel-EDI) on connectivity from the SMG to the IFJ (**Figure 4-C**). This modulation suggests that under moderate blue-enriched light conditions, the influence of the SMG on the IFJ is increased. Low illuminance orange light (0.16 mel-EDI) showed positive evidence (P=0.93) for a strengthening impact on the connectivity from the MDN to the SMG (**Figure 4-C**). This suggests that in the evening, during orange conditions, the influence of the MDN on the SMG increases.

This first analysis established the connectivity patterns in the reference group that were also present in the other 2 groups. We then assessed separately light-induced modulation that came in addition to those shared effects by contrasting the reference group with 1) the morning young adult group and 2) the adolescent evening group to identify differences related to the time of day and age.

### Modulation of Thalamus–Prefrontal Connectivity in the Morning

The DCM analysis comparing the impact of illuminance on connectivity revealed positive evidence (Pp=0.80) that the highest illuminance in the morning (190 mel-EDI) strengthened the connectivity from the MDN to the IFJ, whereas this effect was not observed in the evening (**Figure 4-D**). The strengthened modulation of this thalamo-cortical connectivity suggests that, in addition to the light effects detected in the reference group, blue-enriched light has a time-dependent effect on brain connectivity in young adults, and in the context of executive functioning and in the morning, blue-enriched light may modulate neural processing and connectivity through this pathway more effectively.

### Modulation of Thalamus–Parietal Connectivity in Adolescents

The DCM analysis of the modulatory impact of light illuminance across different age groups indicated, with positive evidence (0.92), that moderate illuminance blue-enriched light strengthened the connectivity from the MDN to the SMG in adolescents in the evening, whereas this effect was not observed in adults (**Figure 4-E**). This suggests developmental differences in the brain’s responsiveness to light, with adolescents showing a strengthened influence of the thalamus on SMG activity under moderate illuminance in the evening, in addition to the light effects on low illuminance orange light detected in the reference group.

Given the difference in chronotype between morning and evening adults, we repeated the analyses including chronotype as a covariate. This inclusion did not influence the findings regarding the modulatory effect of light in either group (**Figure S2**).

### Absence of a Link between Connectivity and Performance in the Morning

In the final step, we further examined whether there was a link between performance and connectivity metrics affected by light in all 3 groups or in the morning vs. the evening and in adults vs. teenagers. We correlated the performance under a given illuminance and a given group with the connectivity value found to be affected by light with positive posterior probability.

Our analysis revealed no significant effects of connectivity, time of day, age group or the interaction between connectivity and time of day/age group on performance (p>0.09) when we focused on common connectivity modulated by light in all groups (e.g., MDN-to-SMG modulated by orange light (**Figure S3-A**). and SMG-to-IFJ modulated by moderate-intensity blue-enriched light) (**Figure S3-B**).

When we compared the impact of MDN-to-IFJ connectivity modulated by high-intensity blue-enriched light on performance between morning and evening adults, we found a significant interaction between connectivity and time of day (F(1,29)=7.76, p=0.009), although the main effects were not significant (p>0.3). Post hoc analysis revealed a positive association between connectivity and performance in the evening group (t=3.18, p=0.004) but no such association in the morning group (p=0.24) (**Figure S3-C**).

Finally, when examining the impact of MDN-to-SMG connectivity modulated by moderate-intensity blue-enriched light on performance across age groups, no significant main effects of connectivity or age group were found (p > 0.12); however, there was a statistical trend (p = 0.08) for an interaction between connectivity and age group. Post hoc analyses revealed a significant positive association between connectivity and performance in adults (t = 2.36, p = 0.02), but no such association in adolescents (p = 0.6) (**Figure S3-D**).

## Discussion

This study aimed to evaluate the impact of light illuminance on ongoing cognitive brain function, particularly on the crosstalk between brain regions during a cognitive challenge. We determined how changes in illuminance affect connectivity within a brain network sustaining an ongoing auditory executive verbal task composed of the left MDN of the thalamus, the SMG in the parietal cortex, and the IFJ in the prefrontal cortex. We further explored separately how time of day, i.e., morning vs. evening in young adults, and age, with a focus on adolescents and young adults in the evening, could affect the interaction of light with the network connectivity. First, we found that in all three groups, moderate blue-enriched light strengthened cortico-cortical connectivity from the SMG to the IFJ in the evening. Unexpectedly, low levels of orange light also exerted an impact on the thalamo-cortical connectivity from the MDN to the SMG. Second, blue-enriched light impacted the network differently in the morning, with the most intense blue-enriched light also affecting the thalamo-cortical connectivity from the MDN to the IFJ in young adults. Third, adolescents responded differently to light in the evening, with moderate illuminance modifying the thalamo-cortical connectivity from the MDN to the SMG. Overall, these results reinforce the view that the thalamus is key in mediating the acute impact of light on cognitive brain function and emphasize that the impact of light depends on the context of light administration. Our findings may have implications for the growing interest in developing lighting interventions aimed at optimizing cognition across ages.

Prior human fMRI research has repeatedly identified the thalamus as the subcortical brain area most consistently affected by light exposure during non-visual cognitive tasks[12,13,15]. Owing to its diverse nuclei, the thalamus plays a crucial role in processing and forwarding sensory information to the cerebral cortex. It further integrates diverse types of inputs critical for cognitive function, including alertness signals, through so-called thalamo-cortical loops. The pulvinar, which occupies a large part of the posterior part of the thalamus, has often been proposed as the thalamic nucleus that mediates light impact, and we recently provided support for this idea in the context of morning light exposure and an auditory attentional task[16]. Focusing on executive functions in the context of an auditory verbal n-back task, our current univariate analyses pointed toward the MDN, which is more anterior than the pulvinar. Importantly, the thalamic ROI we focused on in our prior publication on the impact of light on attention also encompassed smaller portions of several thalamic nuclei in addition to the pulvinar, including the MDN[16]. Similarly, the current thalamus ROI was focused on the MDN but extended dorsally to include a small portion of the pulvinar. Both publications therefore focus on the same brain structures even if their pulvinar vs. MDN gradient is distinct. Like the pulvinar nucleus, the MDN is an associative nucleus (not only relaying sensory information to the cortex), and it is important for executive functions such as attention, working memory and decision making[52–54]. It is therefore in good position to convey the stimulating influence of light to the cortex.

Our univariate analyses further indicated that, similar to the attentional task, the SMG was strongly involved in ongoing processes (the IPS reported in our prior publication consists of the sulcus continuing the gyrus of the SMG), as was the IFJ, which is more specific to executive function. In line with the verbal nature of the task, the responses were stronger over the left hemisphere, so we focus on a network composed of region of that hemisphere. The connectivity between the thalamus and both cortical regions is of particular interest. The SMG plays an important role in working memory, particularly in verbal working memory tasks[18]. As part of the parietal lobe, the SMG is involved in the (mostly left-lateralized) phonological loop[18,55], is essential for storing serial order information and manipulating verbal information and plays a key role in focusing and controlling attention. The connectivity between the thalamus and the SMG, which we detect irrespective of light illuminance, may enhance the ability of the SMG to integrate sensory information and support phonological processing. The IFJ located within the frontal cortex is a key structure for cognitive control, particularly in tasks that demand executive functioning involving decision-making, such as the n-back task[19]. It helps with managing information retrieval and updating. The connectivity between the thalamus and the IFJ, which we also detect irrespective of light illuminance, can facilitate this process by modulating responses on the basis of current cognitive demands and incoming sensory information and ensuring that decision-making and executive processes are informed by the latest sensory inputs.

When considering the impact of changing light illuminance on the 3-ROI network involved in the ongoing task, we must first emphasize that we cannot determine which retinal photoreceptor is responsible for the connectivity changes we observed. While our rationale for conducting the study is based on the maximal sensitivity of ipRGCs to shorter wavelengths, all conditions varied in terms of rod, cone and melanopsin stimulation. The light levels and spectra we administered likely contributed to our findings, with cones potentially contributing through their inputs to ipRGCs[56]. While a classical response on rods is less likely, it cannot be ruled out[57].

The impact of moderate illuminance on cortico-cortical connectivity from the SMG to the IFJ in all participants (with moderate evidence) is in line with the findings of a previous study showing that blue light exposure interacts with the ongoing cognitive process by increasing the functional connectivity between the prefrontal cortex and multiple cortical regions, including the SMG[58]. This likely corresponds to a strengthening impact of top-down attentional processes mediated by the SMG on higher-order cognitive functions, potentially strengthening the integration of attentional processes with executive functions. Additionally, the fact that only moderate illuminance blue-enriched light affected SMG-IFJ connectivity may be due to a ceiling effect, where higher intensities do not yield additional effects or a response that follows an inverted U-shaped function.

Surprisingly, we further found that the connectivity from the MDN to the SMG was influenced by low illuminance orange light, to which ipRGCs should be only weakly sensitive. This likely reflects the involvement of rods and/or cones that trigger a visual response, ultimately affecting the crosstalk between the MDN and SMG. Given that the MDN has been implicated in visual responses, our findings may reflect the influence of visual responses on alertness and attention, reminding that light effects always consist of a mixture of visual and non-visual responses. Similar to the impact of blue-enriched light on SMG-to-IFJ connectivity, the interaction impact of orange light on the ongoing cognition could increase/optimize MDN-to-SMG connectivity and thereby the attentional resources required for ongoing processes.

Rods and cones signals can also reach the posterior thalamus and the SMG through the visual pathway, with relays in the lateral geniculate nucleus (LGN) and in the primary visual cortex. IpRGCs can directly reach the thalamus through the paraventricular nucleus[59] or the ventral LGN (the intergeniculate leaflet in rodents)[60] or directly through the pulvinar[61]. IpRGCs signaling could also indirectly reach our networks through their dense projection to hypothalamus nuclei, including the lateral hypothalamus (LH), as we recently suggested based on part of the same dataset[21], as well as through the locus coeruleus (LC), which receives indirect inputs from the suprachiasmatic nucleus, orchestrating circadian rhythmicity and of the LH. Given the broad projections of the LH and LC and their involvement in alertness regulation, they could influence both MDN and SMG activity and connectivity. Determining which of these options is more likely will require further complex connectivity analyses, including other brain regions, such as the LC and LH.

Importantly, when we focused on time of day differences in young adults, we found that in the morning, the highest illuminance affected the MDN–IFJ connectivity. This could again correspond to the translation of the increase in alertness by light into enhanced frontal activity to improve ongoing high-order executive processes. This is reminiscent of the impact of morning blue-enriched light on thalamus/pulvinar connectivity to the parietal cortex[16]. The interaction impact of morning light onto cognition may therefore primarily affect thalamocortical loops to the parietal cortex in case of attentional tasks and to both the parietal and frontal cortex in more complex executive tasks[62]. This change in the crosstalk from the thalamus to the cortex would come on top of the parietal to prefrontal changes in connectivity, at least for executive processes, as this connectivity was not tested for attentional processes, warranting further investigations. Since the highest illuminance did not affect MDN-to-IFJ connectivity in the evening, our findings suggest that the dynamics of the impact of illuminance and its potential ceiling effect and/or inverted U-shaped pattern vary throughout the day. The previous suggestion that light may be more beneficial in the morning than in the evening, when considering brain function from a regional activation perspective[8], may extend to connectivity, which is influenced by a broader range of illuminance in the morning. Light may be more effective at improving alertness and/or attention when one has been awake and/or exposed to light for only a few hours, compared to the end of the day.

Critically, when comparing age groups in the evening, we found that moderate illuminance increased MDN-to-SMG connectivity in adolescents compared with young adults. These findings suggest that in this age group, evening blue-enriched light increases the impact of the thalamus on the attentional resources sustained by the SMG. This would come on top of the influence of orange light on the MDN-to-SMG connectivity (as well as moderate blue-enriched light on the SMG-to-IFJ connectivity). Adolescents may therefore be more affected by light than young adults in the evening, particularly in terms of connectivity, which may influence attention [14]. This may be important given the current concerns regarding adolescents using LED screen devices, particularly in the evening[63]. This further supports the development of specific light interventions in this age group to optimize evening alertness, attention and sleep in the evening, as investigated by others[64,65].

The age-group difference we detected may, however, also arise from differences in the maturation or wiring of the brain, which is considered to be fully completed by the age of 25 years[66]. The group difference could indeed be present at all times of day, i.e., also in the morning, as it was not assessed in the present study. We further note that the fact whereby only moderate illuminance blue-enriched light affected MDN-SMG connectivity may reflect an inverted-U shape rather than a ceiling effect, as higher intensities did not yield additional effects in either age group. Future studies with more light conditions and comparing age groups at different times of day are needed to test the hypotheses we raised here. These studies should also include larger sample sizes, as despite our relatively large research effort to collect data from more than 50 individuals, the different age groups were still composed of 17 to 20 individuals, which may have hindered statistical power. We further note that although sex was controlled for in the statistical analyses, we have been insufficiently powered to detect sex differences between time of day and/or age groups. Also, although we only focused on the left cortical hemisphere and diencephalon given the verbal nature of the task, we posit that the effects we detected would be generalizable to the right hemisphere within an executive context, e.g. for right-lateralized spatial working memory task. This remains, however, to be demonstrated.

Finally, we refer to recent recommendations for the light level, which should be used during the day (250 mel-EDI lux), in the evening (10 mel-EDI lux) and at night (darkness: 1 mel-EDI lux) to enhance physiology, sleep quality, and wakefulness in healthy adults[3]. The highest illuminance we administered was 190 mel-EDI, which impacted brain connectivity in the morning, confirming that illuminance in a relatively similar range to the recommendation affects brain functions. The moderate illuminance we administered was 92 mel-EDI lux and impacted connectivity in all groups (particularly in adolescents), confirming that higher illuminance than recommended for the evening can affect brain functions.

Concomitant with the changes we reported at the level of the brain, light illuminance affected performance on the task, with better accuracy under higher illuminance. We therefore posit that the changes in connectivity triggered by light underline part of the behavioral changes, but they cannot be attributed to the impact of light on a single connection. As a results, we did not consistently observe a direct link between task accuracy and the connectivity metrics influenced by light. Given the absence of a difference in performance under the highest intensity blue light between morning and evening adults, the link between connectivity and performance under the highest intensity blue light that was found only in the evening group may reflect the role of neurophysiological states in how connectivity translates into behavior. For example, higher levels of alertness or arousal in the morning could make performance less dependent on connectivity strength. Focusing on evening groups, i.e., adolescents and adults with evening fMRI data, an association between connectivity and performance under moderate blue light was found, although it did not reach a significant level. This trend could suggest that the connection between connectivity strength and performance in the evening under lower-intensity light may not be very strong, and a larger sample size might be required to uncover this more clearly. All these behavioral findings remind us that behavior depends on complex interactions between the (cortical) region of a much larger network than the one we focus on in the present study and warrants future, more complex connectivity investigations.

In summary, we show that light influences information flow from the thalamus to cortical areas and between cortical regions, potentially reflecting how the impact of light on alertness and attention facilitates integration in a network highly relevant for executive functions. These effects are not detected in the entire network sustaining an ongoing cognitive process and are specific to the crosstalk between certain brain regions, with a potential central role of the posterior and dorsal thalamus (pulvinar and/or MDN). We further report that the interaction between light and cognition varies both with environmental and individual factors, i.e. with the time of day and brain development stage. Overall, our findings add to the growing body of research building the promise of light interventions to optimize brain functions (and sleep) at different times of day and across the lifespan[1].

## Supporting information

Revised Supplementary Material

## Acknowledgments

The study was conducted at the GIGA-In Vivo Imaging technological platform of ULiège, Belgium. The authors thank Christine Bastin, Alexandre Berger, Christina Schmit, Annick Claes, Christian Degueldre, Catherine Hagelstein, Gregory Hammad, Brigitte Herbillon, Patrick Hawotte, Sophie Laloux, Erik Lambot, Benjamin Lauricella, Pierre Maquet, Eric Salmon and Siya Sherif for their help during the different steps of the study.

## Ethical Approval

The study was approved by the Ethics Committee of the Faculty of Medicine of the University of Liège. All participants provided their written informed consent and received monetary compensation for their participation.

## Author Contributions Statement

R.S. and G.V. designed the research. R.S., F.B., I.C., I.P., and E.B. acquired the data. N.M., E.K., J.R., F.C., C.P., P.T., L.L. and M.Z. provided valuable insights while acquiring, interpreting, and discussing the data. R.S. analyzed the data and was supervised by G.V. R.S. and G.V. wrote the paper. All authors edited and approved the final version of the manuscript.

## Data availability

The processed data and analysis scripts supporting the results included in this manuscript are publicly available via the following open repository: https://gitlab.uliege.be/CyclotronResearchCentre/Public/xxxx (the repository will be created following acceptance/prior to publication of the paper). The raw data could be identified and linked to a single subject and represent a large amount of data. Researchers willing to access the raw data should send a request to the corresponding author (GV). Data sharing will require evaluation of the request by the local Research Ethics Board and the signature of a data transfer agreement (DTA).

