## Supplementary material for "Light Impact on Thalamo-Cortical Connectivity During a Cognitive Task Depends on Time of Day and is Different in Teenagers": Revised Supplementary Material

**Table S1: Characteristics of the light conditions.**

| | Low BEL | Mid BEL | High BEL | Monochromatic Light (589nm) |
|---|---|---|---|---|
| **Illuminance (lux)** | 47 | 116 | 240 | 7.5 |
| **Peak Spectral Irradiance (nm)** | 460 | 460 | 460 | 590 |
| **Melanopic EDI (lux)** | 37 | 92 | 190 | 0.16 |
| **Rhodopic EDI (lux)** | 39 | 97 | 201 | 0.94 |
| **Cyanopic EDI (lux)** | 32 | 79 | 163 | 0 |
| **Chloropic EDI (lux)** | 44 | 110 | 227 | 5 |
| **Erythropic EDI (lux)** | 46 | 113 | 233 | 8 |
| **Irradiance (µw/cm²)** | 15 | 36 | 75 | 1.4 |
| **Photon Flux(1/cm²/s)** | 4.12E+13 | 1.02E+14 | 2.10E+14 | 4.24E+12 |
| **Log Photon Flux ($Log_{10}$ (1/cm²/s)** | 13.61 | 14.01 | 14.32 | 12.63 |
| **Narrowband Peak** | - | - | - | 589 |
| **Narrowband FWHM** | - | - | - | 10 |

Detailed light characteristics of the four light conditions used in fMRI protocol. Blue enriched light (BEL) (low, mid, and high) and monochromatic light (589nm).

**Table S2: Activated brain regions in response to the auditory task (contrast: 2-back – 0-back)**

| **Brain area** | **Hemisphere** | **X,Y,Z** | **P-value (FWE corrected)** |
|---|---|---|---|
| Cerebellum | L | -26, -62, -33 | <0.0001 |
| | R | 30, -63, -29 | <0.0001 |
| Anterior Insula | L | -32, 21, 2 | <0.0001 |
| | R | 35, 23, -1 | <0.0001 |
| **Supramarginal gyrus (SMG)** | L | -38, -46,41 | <0.0001 |
| | R | 42, -42, 45 | <0.0001 |
| **Inferior frontal junction (IFJ)** | L | -47, 10, 32 | <0.0001 |
| | R | 52, 14, 27 | 0.004 |
| **Thalamus (MDN)** | L | -11, -19, 10 | <0.0001 |
| | R | 11, -19, 10 | <0.0001 |
| Precuneus | L | -5, -65, 54 | 0.002 |
| | R | 5, -65, 50 | <0.0001 |
| Superior frontal gyrus | L | -23, 5, 52 | <0.0001 |
| | R | 25, -2, 54 | <0.0001 |
| Caudate | L | -16, -1, 18 | <0.0001 |
| | R | 17, -4, 20 | <0.0001 |
| Putamen | L | -20, 2, 5 | 0.006 |

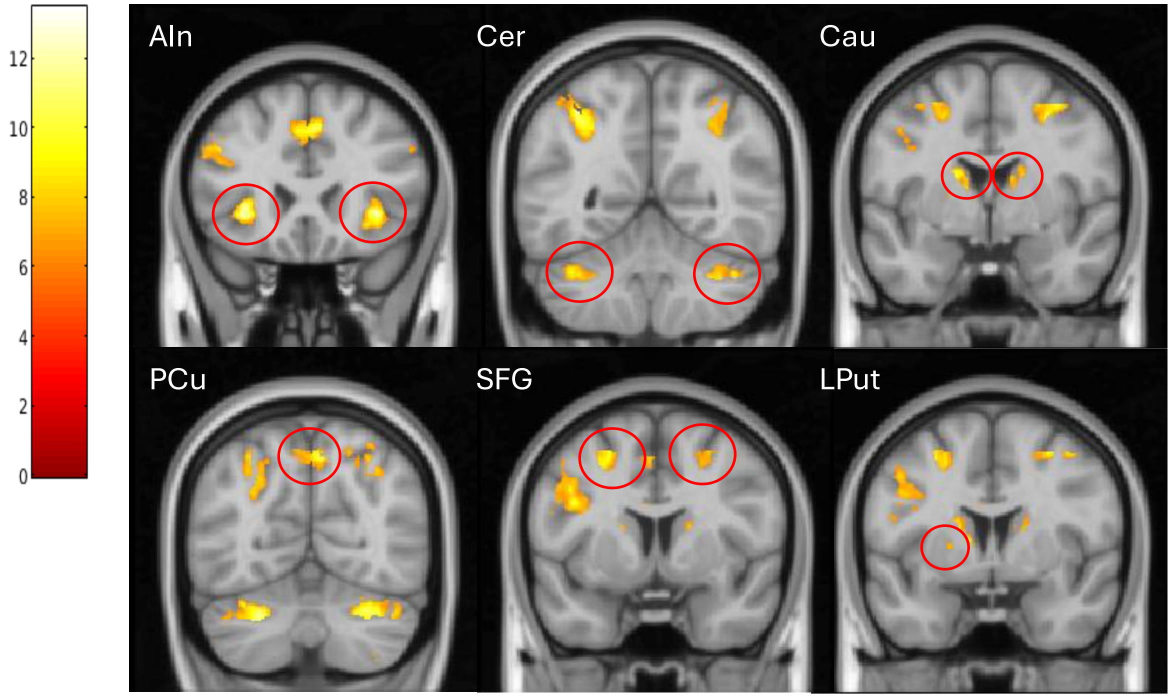


**Figure S1. Brain responses to the task (Contrast: 2-back vs 0-back).** Bilateral brain activations ($P_{FWE}$ = 0.05) in Anterior Insula (AIn), Cerebellum (Cer), Caudate (Cau), Precuneus (PCu), Superior Frontal Gyrus (SFG) and Left Putamen (LPut). Refer to Table S2 for the full list of activations.

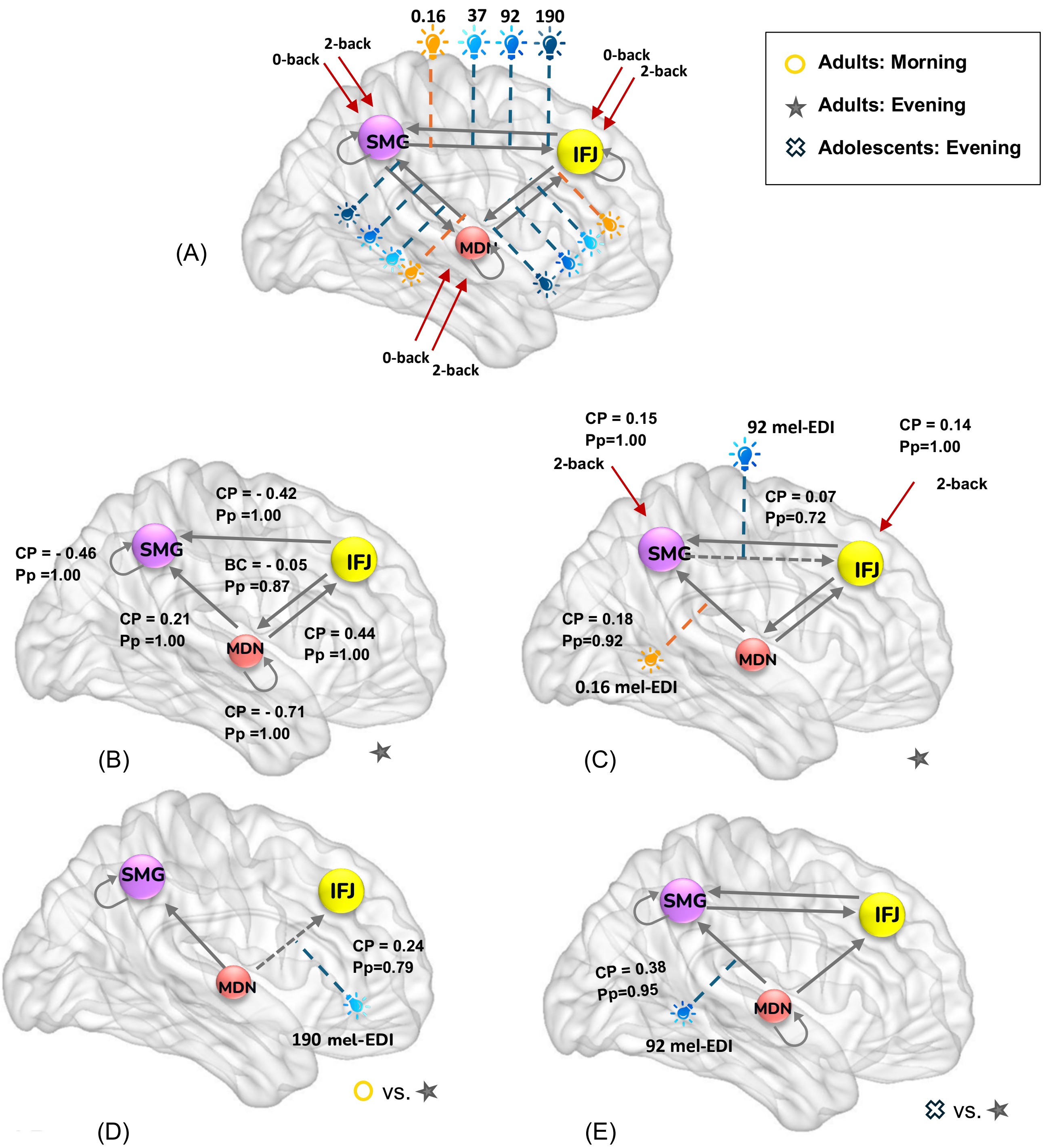


**Figure S2: Effective connectivity results controlling for chronotype: (A)** initial model tested using the DCM framework. **(B)** Baseline (intrinsic) connectivity parameter (CP) common to all groups, as isolated in the reference group (i.e., Adults with fMRI in the evening). **(C)** Modulatory impact of light conditions on connectivity along with task input in the reference group (common to all groups). **(D)** <u>Differences</u> between the connectivity parameters in the morning compared with those in the evening (in adults). **(E)** Differences in the connectivity parameters between adolescents and adults (both in the evening).

IFJ: inferior frontal junction; MDN: mediodorsal nucleus of the thalamus; mel-EDI: melanopic equivalent daytime illuminance. Pp: posterior probability; SMG: supramarginal gyrus; gray arrows: between regions and self-inhibitory connectivity; red arrows: task inputs; colored light bulb: light conditions as connectivity modulators (orange: 0.16 mel-EDI; dark blue: 37 mel-EDI; medium blue: 92 mel-EDI; light blue: 190 mel-EDI).

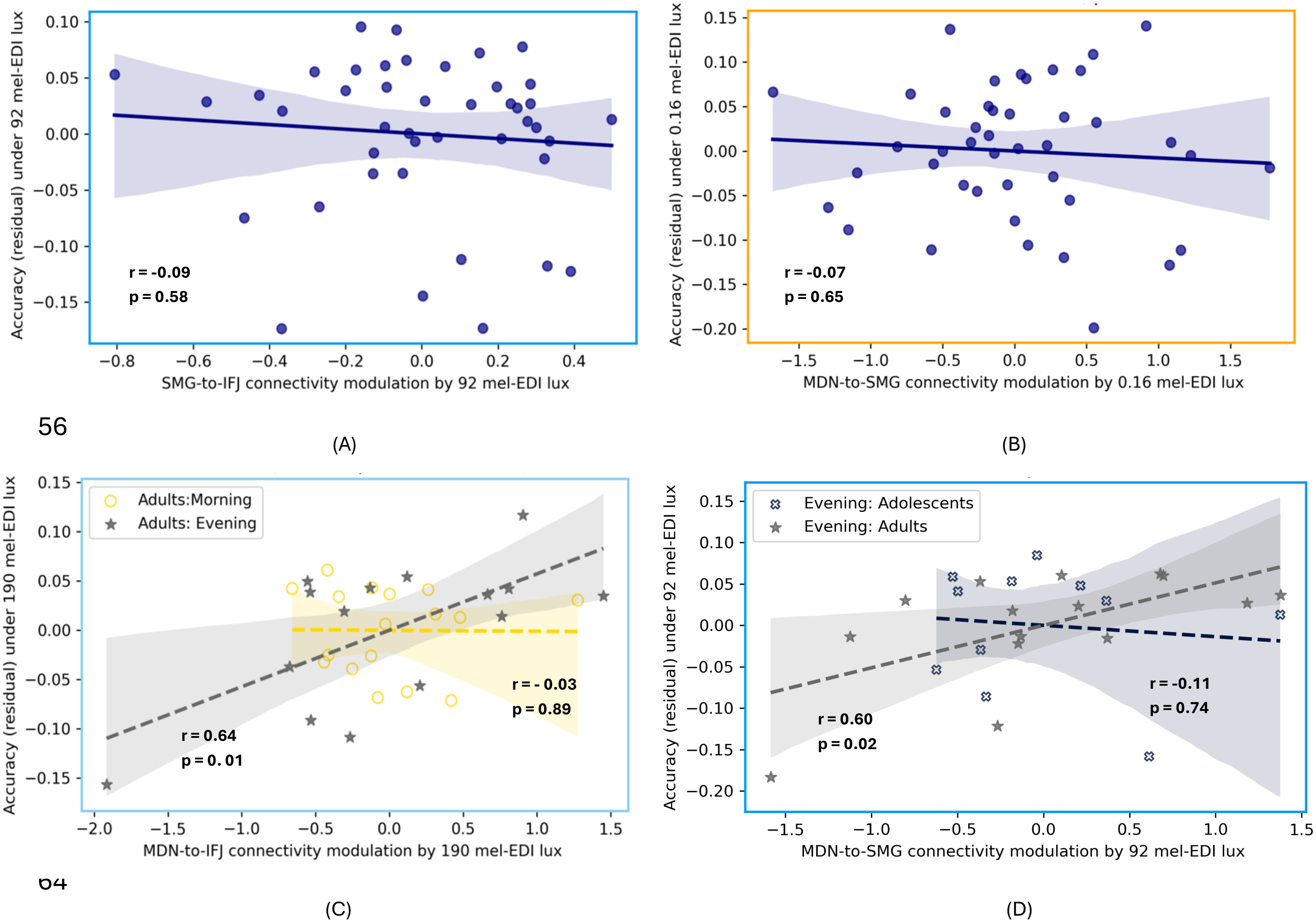


**Figure S3: Linear regression plots showing the relationship between light-modulated connectivity and task performance, controlling for sex, BMI.**

(A) SMG-to-IFJ connectivity under moderate-intensity blue-enriched light and (B) MDN-to-SMG connectivity under orange light showed no significant association with performance when examining all participants together.

(C) MDN-to-IFJ connectivity under high-intensity blue-enriched light showed a significant interaction with time of day, with a positive association in evening adults (r = 0.64, p = 0.01).

(D) MDN-to-SMG connectivity under moderate-intensity blue-enriched light, showed a statistical trend for the interaction with age group with a positive association in adults (r = 0.60, p = 0.02)
